# Differential impacts of drought and esca expression on Ascomycota fungi in the trunks and young organs of mature grapevines

**DOI:** 10.1101/2025.07.03.663063

**Authors:** Pierre Gastou, Giovanni Bortolami, Nathalie Ferrer, Gregory A. Gambetta, Samuele Moretti, Jessica Vallance, Chloé E.L. Delmas

## Abstract

Perennial plant decline is increasingly threatening the profitability and sustainability of agriculture and forestry worldwide. It results from intricate interactions between microbial communities, the plant host, and abiotic stressors. We investigated the effects of drought and esca disease on mature grapevine phytobiomes. Grapevines display no esca leaf symptoms during droughts, but the impacts of drought and esca expression on fungal communities and wood health in mature plants remain poorly understood.

We studied 43 uprooted 30-year-old naturally infected vines in three experimental conditions: well-watered asymptomatic (Control) vines, vines with esca symptoms (Esca), and vines subjected to water deficit (WD) over two consecutive summers. We profiled trunk, cane, stem and petiole Ascomycota communities by DNA metabarcoding with primers specifically designed for grapevine trunk-associated Ascomycota, and quantified wood necrosis. The Ascomycota communities of trunks and younger organs clearly differed, and drought and esca had different impacts on the Ascomycota communities of perennial and young organs. In the trunk, drought significantly decreased fungal diversity in healthy wood and increased the abundance of wood pathogens (e.g. *Phaeomoniella chlamydospora, Botryosphaeria dothidea*). In young organs, esca expression decreased the species richness and diversity of the Ascomycota community to a greater extent than drought. We also found that the relative proportion of healthy wood was smaller in plants with esca symptoms than in control plants.

Thus, drought increased Ascomycota pathogen abundance in the trunk but did not increase wood degradation and esca expression, highlighting the need to investigate the molecular basis of plant-microbiome interactions under multi-stress conditions.

## Introduction

A balanced plant microbiota is essential to preserve plant health and productivity in a context of global changes and emerging biotic pressures (Compant et al. 2019; Trivedi et al. 2020, 2022). The disequilibrium or dysbiosis of endophytic microbiota has been associated with plant dieback, although the causal mechanism is generally unknown (Arnault et al. 2023, Bettenfeld et al. 2020, Broberg et al. 2018, Griffiths et al. 2020). Pathogenic and saprotrophic species that are present in a latent endophytic phase within healthy plants (Hallmann et al. 1997, Marsberg et al. 2017) are often overabundant or more active in symptomatic plants, potentially worsening the decline of these plants (Bettenfeld et al. 2020, Jiao et al. 2023).

Plant microbial communities are also affected by abiotic factors, and these interactions are associated with plant dieback events worldwide (Ciesla and Donaubauer 1994, Desprez-Loustau et al. 2006, Manion 1981). In a context of climate change, the effects of drought on interactions between plant physiology, plant pathogens and the whole microbial community (i.e. holobiont) are a major issue that cannot be reduced to unequivocal relationships. Depending on plant host species and physiological status, pathogen lifestyle, and the intensity and timing of the stress, drought may exacerbate, inhibit or have no effect on plant-pathogen interactions (Desprez-Loustau et al. 2006, Jactel et al. 2012, Luna et al. 2025, Ramegowda and Senthil-Kumar 2015, Torres-Ruiz et al. 2024). The impact of drought stress on phytobiomes has been documented principally for the rhizosphere and the phyllosphere (Bechtold et al. 2021, Fitzpatrick et al. 2018, Jaeger et al. 2024, Xu et al. 2018) but remains largely unknown for other compartments (e.g. endosphere or woody perennial organs). A comprehensive understanding of the interactions between perennial plant phytobiomes, abiotic and biotic stresses would contribute to deciphering the relative roles of the various factors in plant decline.

Grapevine (*Vitis vinifera* L.), a major cultivated perennial crop, is particularly severely affected by complex dieback processes, mostly involving trunk diseases and environmental factors (Claverie et al. 2020, Etienne et al. 2024). Esca is a trunk disease causing grapevine dieback worldwide, with harmful consequences for the productivity and perenniality of the vineyard (Gramaje et al. 2018). Esca triggers physiological modifications in the leaf, stem and whole plant, impairing growth, hydraulic conductivity, gas exchange, and metabolism within perennial and vegetative organs (Bortolami et al. 2019, 2021a,b, 2023, Chambard et al. 2025, Dell’Acqua et al. 2024a, Magnin-Robert et al. 2016, 2017). This disease is detected in the summer on plant canopies displaying typical symptoms of scorching between the veins of the leaf (so-called “tiger-stripe” symptoms) and apoplexy, which may be observed on some or all of the stems of plants randomly distributed over the vineyard (Lecomte et al. 2024). Internal wood necrosis, with black necrotic tissue and white rot, is observed in both leaf-symptomatic and asymptomatic plants (Bertsch et al. 2013). However, white rot tends to be more frequent in symptomatic plants (Bruez et al. 2020, Maher et al. 2012). The trunk microbiota differs between necrotic and non-necrotic (healthy) wood tissues (Bruez et al. 2020, Elena et al. 2018). In particular, black necroses are characterized by a high abundance of Ascomycota whereas white rot necrosis is caused by Basidiomycota fungi, such as *Fomitiporia mediterranea* (Bruez et al. 2020, Moretti et al. 2021). Conversely, there is no clear difference between the microbial communities of healthy wood collected from plants with and without esca leaf symptoms (Del Frari et al. 2019, Dell’Acqua et al. 2025, Hofstetter et al. 2012, Monod et al. 2024). Several environmental factors are typically involved in perennial plant dieback but drought has a particularly strong impact in viticulture, and this impact is growing even stronger with the acceleration of climate change (van Leeuwen et al. 2024). In this context, the impact of drought on esca pathogens is a growing matter of concern, as both drought and esca have deleterious effects on grapevine hydraulic function.

Recent advances have improved our understanding of the environmental drivers of esca symptom expression, highlighting the key role of drought through its impact on plant water status. Contrasting with the many observations of synergistic impacts of drought and pathogens in trees (Desprez-Loustau et al. 2006, Gomez-Gallego et al. 2022, Torres-Ruiz et al. 2024), esca and drought are antagonist in grapevine, inhibiting each other (Bortolami et al. 2021b; Dell’Acqua et al. 2025). Bortolami et al. (2021b) demonstrated that uprooted naturally infected plants subjected to moderate drought (predawn water potential around -1 MPa) did not express esca leaf symptoms at any time in the course of a two-year experiment. Dell’Acqua et al. (2025) showed that vines with esca symptoms are less affected by intense drought stress and recover more quickly after such stress. The characteristics of fungal communities probably play a key role in drought-esca relationships, but this role has never been elucidated in mature plants.

Drought modifies the structure and functioning of plant microbial communities across compartments (see for example Carbone et al. 2021, Debray et al. 2022, Lin et al. 2023, Trivedi et al. 2022). A recent pluriannual study on young potted vines revealed a significant shift in wood fungal communities during drought, with a decrease in the complexity of microbiota structure and interactions, and an increase in the abundance of *Phaeomoniella chlamydospora*, a pathogenic Ascomycota involved in Petri disease that has been implicated in the esca pathosystem (Leal et al. 2024). These results seem to contradict the antagonism between drought and esca. There is therefore a need to validate these findings on mature grapevines and to consider again which Ascomycota pathogens are associated with the different disease symptoms.

Here, we studied the effects of abiotic (drought) and biotic (esca) stresses on Ascomycota communities in the various organs of grapevine. We addressed the following questions: (i) What impact do seasonal drought and esca leaf symptom expression have on the Ascomycota communities in trunk and younger vegetative organs? (ii) Is the antagonistic relationship between esca and drought associated with shifts in the functional properties of Ascomycota communities? (iii) How do drought and esca expression affect internal wood necrosis and the corresponding Ascomycota communities? We used the same 43 uprooted vines with different esca histories and exposure to contrasting water conditions as described by Bortolami et al. (2021b). We sampled the trunk (separating the necrotic and healthy wood) and young organs (two-year-old canes, current-year stems, and petioles), and used a metabarcoding approach to study grapevine trunk-associated ascomycete fungi.

## Materials and methods

### Plant material and sampling

Sampling was conducted between October 7 and 14 2019 on a subset of uprooted vines (*V. vinifera* L. cv. Sauvignon blanc; *n*=43) from the experimental design described in detail by Bortolami et al. (2021b). This experimental design included three factors: (i) two levels of watering regime in two consecutive summer seasons in 2018 and 2019: WW (well-watered for two years, Ψ_PD_ close to 0 MPa), and WD (water-deficit, presenting Ψ_PD_ ≈ -1 MPa from July to October in 2018 and 2019); (ii) two contrasting histories of esca symptoms since 2012: pA (previously asymptomatic, i.e. plants that had never expressed esca leaf symptoms at any time since 2012), and pS (previously symptomatic, i.e. plants that had expressed esca leaf symptoms at least once since 2012); iii) two canopy phenotypes in 2019: presence of esca leaf symptoms (typical tiger-stripe symptoms) and asymptomatic plants (hereafter referred to as control plants). Plants subjected to a WD never expressed esca leaf symptoms (see Bortolami et al. 2021b). We therefore sampled vines from four distinct groups (Table 1): Water Deficit-pS (WD, *n*=12 asymptomatic plants in 2019); Esca-pS (Esca, *n*=10 well-watered plants expressing tiger-stripe symptoms in 2019); Control-pA (*n*=10, well-watered plants asymptomatic since 2012); Control-pS (*n*=11, well-watered plants asymptomatic in 2019). Esca history had no significant effect (see the Results section), so the Control-pA and Control-pS groups were merged into a single control category (*n*=21, well-watered plants asymptomatic in 2019) for some analyses. During the growing seasons, the plants were regularly treated for powdery mildew (with 80% micronized sulfur-based multisite products).

**Table 1.**
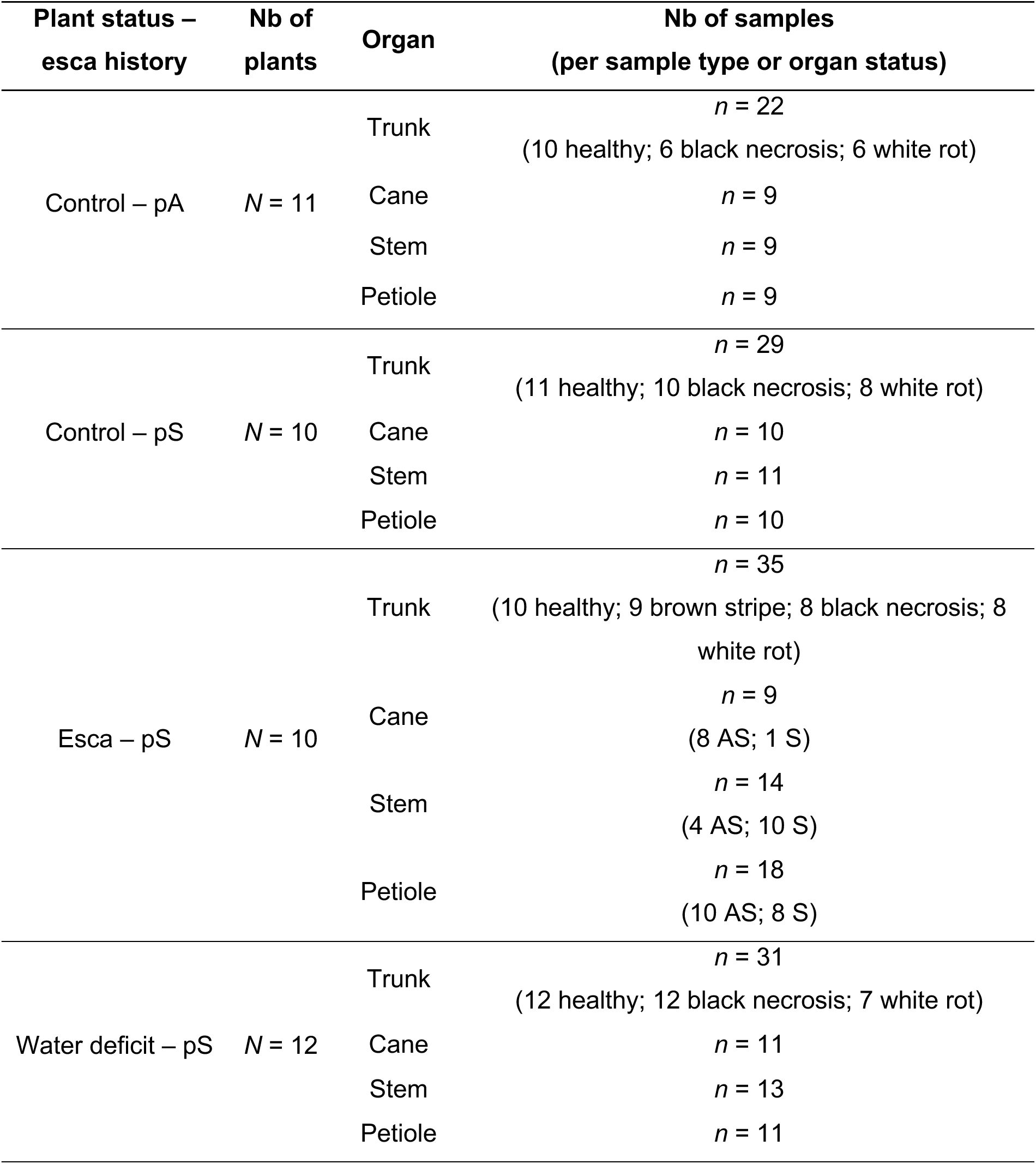
Number of *Vitis vinifera* cv. Sauvignon blanc plants and samples included in this study, classified by experimental plant status, organ and sample types. For esca plants, samples of petioles, current-year stems, and two-year-old canes were classified by organ status: asymptomatic (AS) or presenting tiger-stripe symptoms on the leaves (S). Plants were classified by esca history: previously asymptomatic (pA) or previously symptomatic (pS).

| Plant status –<br>esca history | Nb of<br>plants | Organ | Nb of samples<br>(per sample type or organ status) |
| --- | --- | --- | --- |
| Control – pA | N = 11 | Trunk | n = 22<br>(10 healthy; 6 black necrosis; 6 white rot) |
|  |  | Cane | n = 9 |
|  |  | Stem | n = 9 |
|  |  | Petiole | n = 9 |
| Control – pS | N = 10 | Trunk | n = 29<br>(11 healthy; 10 black necrosis; 8 white rot) |
|  |  | Cane | n = 10 |
|  |  | Stem | n = 11 |
|  |  | Petiole | n = 10 |
| Esca – pS | N = 10 | Trunk | n = 35<br>(10 healthy; 9 brown stripe; 8 black necrosis; 8 white rot) |
|  |  | Cane | n = 9<br>(8 AS; 1 S) |
|  |  | Stem | n = 14<br>(4 AS; 10 S) |
|  |  | Petiole | n = 18<br>(10 AS; 8 S) |
| Water deficit – pS | N = 12 | Trunk | n = 31<br>(12 healthy; 12 black necrosis; 7 white rot) |
|  |  | Cane | n = 11 |
|  |  | Stem | n = 13 |
|  |  | Petiole | n = 11 |

We sampled four different organs from each plant for DNA metabarcoding (Table 1; Fig. S1): petioles (several petioles pooled together), stems from the current year (one debarked internode from the middle of the stem), two-year-old canes (one debarked internode), and trunks (with the separation of necrotic and healthy wood). Consequently, both endophytic and epiphytic Ascomycota communities were included in petiole analyses but only endophytic Ascomycota communities were included in the analyses of woody organs (i.e. stems, canes, trunks). For plants expressing esca symptoms, we separated petioles, stems and canes according to their disease status: asymptomatic (AS) or displaying tiger-stripe symptoms on the leaves (S) (see pictures of leaves in Fig. S1).

We used sterilized tools to remove the bark from the trunk of all plants. We searched for the presence of a brown stripe, a colored band of xylem band appearing under the bark concomitantly with esca leaf symptoms (Lecomte et al. 2012, 2024). A brown stripe was observed in all the plants presenting leaf symptoms (Fig S1) but none of the control plants. We sampled chips from the brown stripe at different heights along the trunk to obtain sufficient material. We then cut a 2 cm-thick cross-section of the trunk 15 cm from the top of the plant and photographed it for further analyses (Fig. S1). We collected tissue of different wood types from this cross-section (after removal of the bark) with sterile shears in a sterile field. This tissue was immediately immersed in liquid nitrogen and stored at -80°C. We collected the following wood types (Table 1): apparently healthy wood, black necrotic wood, white rot, and brown stripe. For white rot, if insufficient necrotic tissue was collected for pathogen detection, we sampled additional white rot tissues from just above or below the cut section. Necrotic tissue was sampled only when present, whereas healthy wood was sampled from every plant. We obtained a total of 251 samples (Table 1): 134 young organs (48 petioles, 47 stems, 39 canes) and 117 trunks (43 healthy wood, 36 black necrosis, 29 white rot, and 9 brown stripes).

### DNA extraction and sequencing

Samples (*n*=251) were ground in liquid nitrogen with a one-ball mill TissueLyser II (Qiagen, Hilden, Germany), taking care to sterilize the balls and sample holders with ethanol 70% [v/v] between samples. Genomic DNA was extracted from 60 mg tissue samples with the Indvisorb Spin Plant mini Kit (Invitek Diagnostics, Berlin, Germany) according to the manufacturer’s instructions. Empty tubes — containing only the extraction reagents, without sample — were included as negative controls for DNA extraction (*n* = 5). DNA was then quantified with a DS-11 spectrophotometer (DeNovix, Wilmington, USA) and adjusted to a uniform concentration of 15 ng x µl^-1^. The ITS2 region of the nuclear ribosomal repeat unit was then amplified with the GTAAf/GTTAr primers (AAAACTTTCAACAACGGATC/GACCTCGGATCAGGTAGGRA), which were specifically designed and optimized for the profiling of pathogenic Ascomycota fungal communities associated with trunk diseases (Morales-Cruz et al. 2018b). DNA was amplified by PCR with an Epgradient Mastercycler (Eppendorf, Hambourg, Germany) in a reaction mixture (25 µl final volume) consisting of 1 µl DNA template (15 ng x µl-1), 10 µl of 2X Platinum Hot Start PCR Master Mix (Invitrogen,Waltham, USA), 0.5 µl of each primer (10 µM), 1.25 µl 20X bovine serum albumin (New England BioLabs, Ipswich, USA), and 11.75 µl DNAse/RNAse-free sterile water. The cycling parameters were as follows: enzyme activation at 95°C for 2 min; 35 cycles of denaturation at 95°C for 45 s, 60°C for 1 min, 72°C for 1.5 minutes; and a final extension at 72°C for 10 min. Empty wells on the PCR plate — i.e. those containing the PCR mix but without template — were used as negative controls (*n* = 9). The PCR products were visualized by electrophoresis in a 2% agarose gel in TBE (molecular weight ≈350 bp). They were then sent to the PGTB sequencing facility (Genome Transcriptome Facility of Bordeaux, Pierroton, France) for sequencing on an Illumina MiSeq platform (v3 chemistry, 2x300 bp). PCR product purification, the addition of multiplex identifiers and sequencing adapters, library sequencing and sequence demultiplexing (with exact index search) were performed by the sequencing service, who included another five negative PCR control samples at this stage.

### Sequence analysis and OTU table construction

Metabarcoding data were processed with the FROGS pipeline (Find Rapidly OTUs with Galaxy Solution) implemented on a Galaxy instance (https://vm-galaxy-prod.toulouse.inra.fr/galaxy/; Escudié et al. 2018). In brief, VSearch was used to merge paired reads with a maximum of mismatch of 10% in the overlapping region. After denoising (i.e. reads with no expected length - between 150 and 590 bp - and containing ambiguous bases, N, were discarded) and the removal of primers/adapters removal with CUTADAPT (Martin 2011), clustering was performed with SWARM (Mahé et al. 2014), with an aggregation distance *d* = 1. Chimeras were removed and low-abundance sequences were filtered out at 0.00005% (i.e. we retained OTUs accounting for at least 0.00005% of all sequences, adapted from Bokulich et al. 2013), to discard singletons from the datasets. The ITS2 region was extracted with ITSx (Bengtsson-Palme et al. 2013) before determining the taxonomic affiliation of the fungal OTUs by BLAST queries against the UNITE Fungi v8.2 (implemented in FROGS), trunkdiseaseID.org (http://www.grapeipm.org/), and NCBI (nr/nt) nucleotide collection (https://blast.ncbi.nlm.nih.gov/Blast.cgi) databases.

We performed a decontamination step, based on 19 DNA extraction and PCR blanks, with the *metabaR v.1.0.0* package (Zinger et al. 2021) implemented in R v.4.2.1 software. Graphical and statistical methods were used to flag and selectively remove non-Ascomycota, putatively contaminating or artifactual OTUs, and potentially unusable samples (i.e. highly contaminated or low sequencing depth).

### Metabarcoding statistical analysis

We compared the diversity, structure and composition of Ascomycota communities (see metrics below) between different categories (Table 1): (i) organs (trunk, cane, stem, petiole; n = 251 samples), considering plant status as a covariate; (ii) plant statuses in healthy trunk wood samples; n = 43); (iii) plant statuses and organ statuses in young organs (n = 134); (iv) trunk wood types (i.e. healthy wood, black necrosis, white rot, brown stripe; n = 117), considering plant status as a covariate; (v) esca symptom histories (i.e. previously asymptomatic *vs*. previously symptomatic) in control plants, considering plant status as a covariate. The (v) analysis was performed separately on trunk (n = 51) and young organ (n = 58) samples. For all analyses, we checked the normality of the residuals (QQ-plot) and the homogeneity of the residual variance (residuals vs. fitted, residuals vs. predictors) graphically. Non-parametric analyses were performed if these assumptions were not satisfied. All analyses were performed with R v.4.2.1 and with an alpha risk of 5%.

To estimate the diversity, structure and composition of Ascomycota communities we calculated three alpha-diversity indices (i.e. observed richness, Shannon index, Simpson index) with the “estimate_richness” function in *phyloseq v.1.42.0* package (McMurdie and Holmes 2013), and compared them across categories by ANOVA procedures followed by Tukey’s post-hoc tests. For Simpson index, the normality assumptions were not fulfilled, so we performed non-parametric alternatives (i.e. Kruskal-Wallis test followed by Dunn’s test). We subjected the Ascomycota community structure of the samples to principal component analysis (PCA) based on a redundancy dependence analysis (RDA) ordination on centered log-ratio (CLR, recommended for compositional data; Gloor et al. 2017)-transformed data, with the *phyloseq* “ordinate” function. We performed a permutational analysis of variance (PERMANOVA) on CLR-transformed data, with the “adonis2” function in the *vegan v.2.6-4* package (Oksanen et al. 2022), to assess the effect of organ, plant statuses, or trunk wood type on Ascomycota community structure, with plant status included as a covariate. The multivariate homogeneity of group dispersions was checked *a posteriori*, and multilevel pairwise comparisons were calculated with the “pairwise.adonis2” function in *pairwiseAdonis v.0.4.1* (Arbizu 2025). A pseudocount of one was added to all OTU counts to handle zero values before CLR transformation, thereby ensuring numerical stability during log-ratio calculations.

For some analyses, we classified Ascomycota OTUs into three groups according to function: putative grapevine wood pathogens, putative antagonists of grapevine wood pathogens (tested with success *in vitro*, in the greenhouse or in the vineyard), and other taxa. This taxonomic classification to genus level was based on the available scientific literature (Supplementary Table S1). The effect of plant status, or wood type on these functional classes was evaluated by ANOVA followed by Tukey’s post-hoc tests.

We performed differential analysis with linear discriminant analysis effect size (LEfSe) to identify the Ascomycota species most likely to explain differences between wood types, with the “run_lefse” function in the *microbiomeMarker v.1.4.0* package (Cao et al. 2022) and the following parameters: kw_cutoff = 0.05 and lda_cutoff = 2.

We performed a differential analysis with the Aldex2 procedure to identify the Ascomycota species most likely to explain differences between control and Esca plants, and between control and WD plants. In this case, the “run_aldex” function was applied to RLE-transformed data, with a *p*-value cutoff of 0.05.

### Analysis of trunk necrosis

We used ImageJ software (Schneider et al. 2012) to analyze each trunk cross-section image. We measured the area of apparently healthy wood, black necrosis, and white rot (in mm²) relative to the entire cross section. The brown stripe was not visible on the cross sections because it affects only the outermost xylem rings and was not, therefore, quantified as for the other tissues.

We assessed the effect of plant status (control, esca, WD) on the relative proportions of healthy wood, black necrosis and white rot in trunk cross sections (n = 43). The effect of plant status on the statistical distribution of wood type proportions was assessed with Fisher’s exact test (simulated *p*-value, 2,000 replicates). The effect of plant status on the proportion of each type of necrosis was assessed by ANOVA followed by Tukey’s post-hoc tests with “HSD.test” in the *agricolae v.1.3-7* package (de Mendiburu 2023). We then plotted the statistical distribution of healthy wood and compared it across plant statuses, using the minimal proportion of healthy wood in control plants as a threshold. Finally, we assessed the effect of esca symptom history on the proportion of each type of necrosis in control plants (n = 21).

## Results

### General description of the Ascomycota community and differences between organs

In total, 10,765,305 reads were generated and assigned to 320 operational taxonomic units (OTUs) (Supplementary Tables S2 and S3). The final decontaminated dataset contained 9,641,320 reads (i.e. 10.4% of reads were removed) from 251 plant samples that were affiliated to 297 OTUs (i.e. 7.2% were removed). Ascomycota communities were composed of four main classes: Dothideomycetes (59.5%), Eurotiomycetes (19.0%), Sordariomycetes (18.4%) and Leotiomycetes (2.2%). Less than 1% of OTUs were affiliated to multiple classes or to no class. Global metrics for both the raw and decontaminated datasets are provided in Supplementary Table S2. The decontaminated OTUs are presented in Supplementary Table S3. The number of reads per sample after decontamination and sample metadata are provided in Supplementary Table S4.

We compared the diversity and structure of Ascomycota communities between plant organs (i.e. trunk, cane, stem, petiole) across all samples. Shannon and Simpson indices differed significantly between organs, whereas observed richness did not (*P* = 0.005, *P* = 10^-4^ and *P* = 0.10, respectively). Ascomycota richness and diversity were slightly lower in trunk samples than in other organs (Supplementary Fig. S2A). PCA and PERMANOVA analyses revealed that organ accounted for 21% of the variance of the total Ascomycota community (*P* = 0.001). Trunk samples were clearly separated from canes, stems and petioles (Supplementary Fig. S2B). We therefore decided to compare trunk samples and young organ samples (canes, stems and petioles, corresponding to organs that were no more than two years old) in subsequent analyses.

### Effects of drought and esca on the Ascomycota community in healthy trunk wood samples

Three species predominated in healthy trunk wood samples: *Phaeomoniella chlamydospora* (28.2%), *Cladosporium sphaerospermum* (13.0%) and *Phaeoacremonium minimum* (8.1%; Fig. 1A). Plant status significantly influenced the Shannon and Simpson indices of healthy wood but not observed richness (*P* = 0.01, *P* = 0.04 and *P* = 0.17, respectively). Ascomycota diversity was significantly lower in the healthy wood of WD plants than in that of control plants, whereas it did not differ significantly between esca and the other two experimental conditions (Fig. 1B). PCA and PERMANOVA analyses revealed that plant status accounted for 6.9% of the variance of the Ascomycota community in healthy wood (*P* = 0.001). WD plants differed significantly from control plants, whereas esca plants were intermediate (Fig. 1C).

**Fig. 1.**
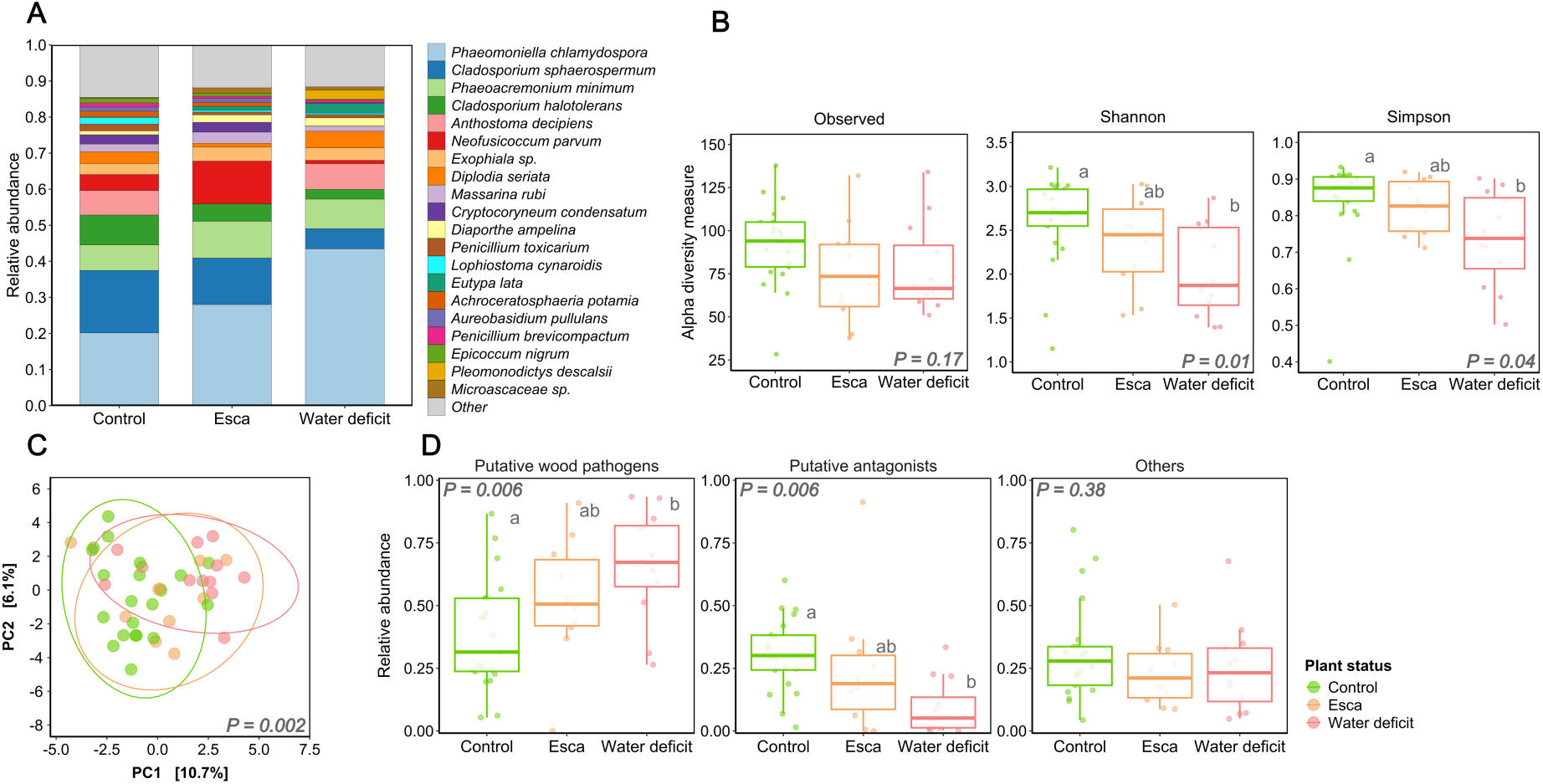
Properties of endophytic Ascomycota communities in healthy trunk samples according to plant status. **A,** Distribution of the 20 most abundant Ascomycota OTUs in trunk samples, by plant status. The mean relative abundance was calculated for each OTU, by plant status. *Stacked bars are colored according to the OTU. Less abundant OTUs are grouped in the category “Other” and are colored in light gray.* **B**, Effect of plant status on alpha-diversity metrics (i.e. observed richness, Shannon and Simpson indices) associated with Ascomycota communities. **B**, PCA summarizing the structure of Ascomycota communities according to an RDA on CLR-transformed data. Ellipses correspond to the 95% interval for each group. **C**, Relative abundance of putative wood pathogens, putative antagonists and other taxa, by plant status. All graphs are colored according to plant status.

Plant status significantly affected the relative abundance of putative grapevine wood pathogens and antagonists (Supplementary Table S1) in healthy wood, but did not affect the abundance of other taxa (*P* = 0.006, *P* = 0.006 and *P* = 0.38, respectively; Fig. 1D). The relative abundance of putative pathogens was significantly higher in WD plants than in control plants (Fig. 1D), whereas the abundance of putative antagonists was significantly lower in WD plants than in control plants (Fig. 1D).

Finally, Aldex2 differential analyses to identify Ascomycota species highlighted differences in microbial composition between plants with different disease statuses. One phylum-affiliated Ascomycota OTU was significantly less abundant in plants with esca symptoms than in control plants. Four fungal species, including the pathogens *P. chlamydospora* and *Botryosphaeria dothidea*, were significantly more abundant in WD plants than in control plants. Conversely, four fungal species, including putative antagonist *Cladosporium* spp. and *Epicoccum nigrum*, were significantly less abundant in WD plants than in control plants (Fig. 1A; Supplementary Table S5).

### Effects of drought and esca on Ascomycota communities in young organs

We first looked for differences in the Ascomycota community between asymptomatic and esca-symptomatic young organs from plants with esca symptoms. We found no significant difference between these two groups of organs for either alpha-diversity or beta-diversity metrics (Supplementary Table S6).

We then compared the Ascomycota communities in young organs between control, esca (pooling samples from asymptomatic and symptomatic canes, stems and petioles) and WD plants. Three species predominated in young organs: *Cladosporium sphaerospermum* (44.5%), *Cladosporium halotolerans* (20.0%), and *Diaporthe ampelina* (5.1%; Fig. 2A). The genus *Cladosporium* accounted for 69.1% of the OTUs retrieved from young organs.

**Fig. 2.**
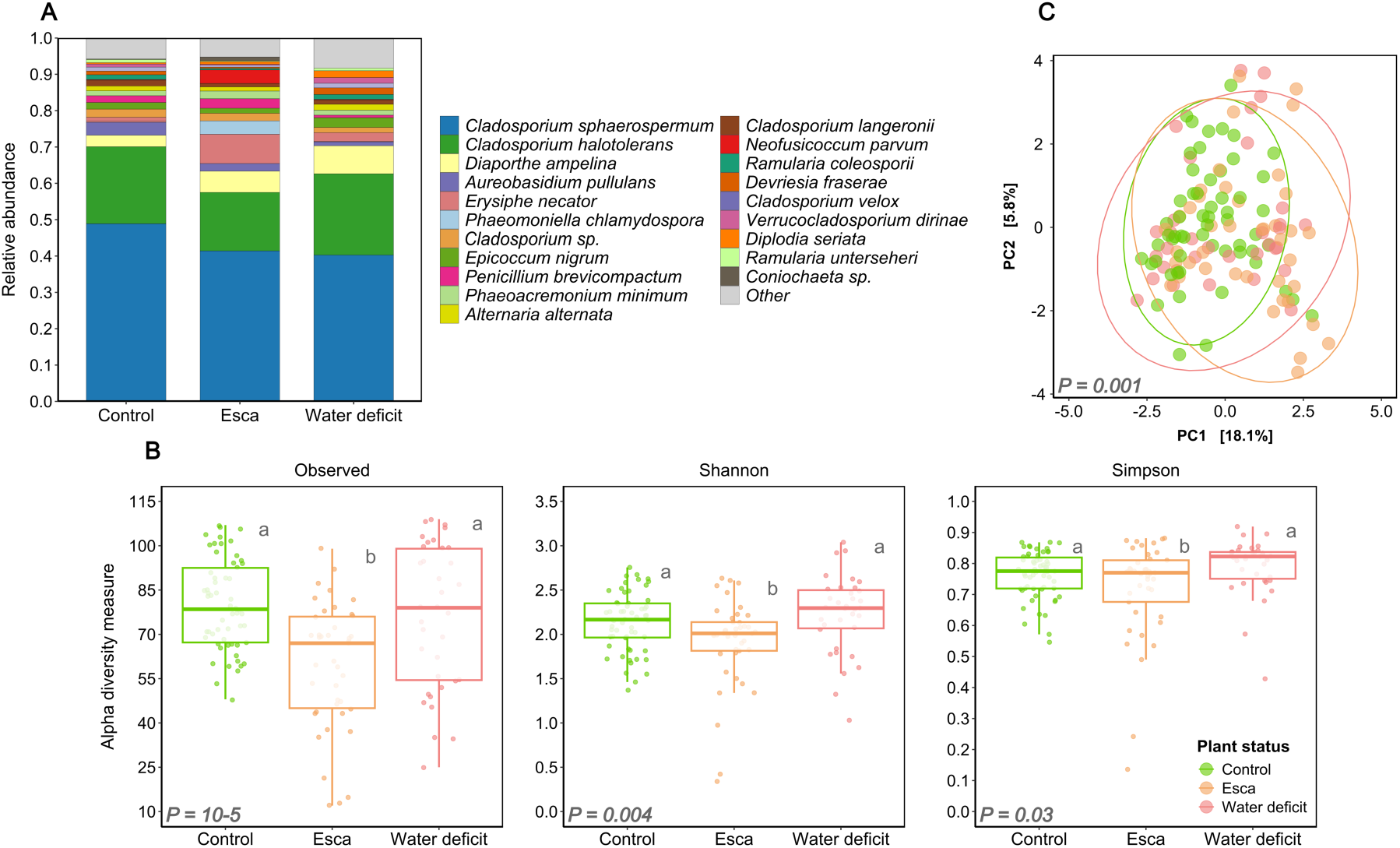
Properties of endophytic and epiphytic Ascomycota communities in young organs (i.e. petioles, stems and canes), by plant status. A, Distribution of the 20 most abundant Ascomycota OTUs, by plant status, in young samples. Mean relative abundances were calculated for each OTU, by plant status. *Stacked bars are colored according to the OTU. Less abundant OTUs are grouped into the “Other” category and colored in light gray.* B, Effect of plant status on alpha-diversity metrics (i.e. observed richness, Shannon and Simpson indices) associated with Ascomycota communities. C, PCA summarizing the structure of Ascomycota communities according to an RDA on CLR-transformed data. Ellipses correspond to the 95% interval for each group. Panels B and C are colored according to plant status.

Plant status significantly influenced observed richness, and the Shannon and Simpson indices (*P* = 10^-6^, *P* = 0.003 and *P* = 0.02, respectively). Ascomycota diversity was lower in the young organs of esca plants than in those of control and WD plants (Fig. 2B). PCA and PERMANOVA analyses revealed that plant status accounted for 5.5% of the variance of the Ascomycota community in young organs (*P* = 0.001). In pairwise comparisons, there were slight differences between plant statuses (Fig. 2C).

We performed Aldex2 differential analyses to identify the Ascomycota species underlying the differences in microbial composition between the young organs of plants with different disease statuses. *Erysiphe necator* was significantly more abundant in esca plants than in control plants. Conversely, 12 fungal species, including several putative antagonist *Cladosporium* spp. and *Aureobasidium pullulans*, were significantly less abundant in esca plants than in control plants. Two fungal species, *Exophiala* sp. and *B. dothidea*, were found to be significantly more abundant in young organs from WD plants than in those of control plants. Conversely, six fungal species, including several *Cladosporium* spp. and *A. pullulans*, were found to be significantly less abundant in WD plants than in control plants (Fig. 2A; Supplementary Table S5). Overall, putative wood pathogens accounted for 11.8 ± 0.2% of the OTUs in young organs. These pathogens were slightly more abundant in esca and WD plants than in control plants (*P* = 0.05). Putative antagonistic genera accounted for 72.9 ± 0.2% of OTUs in young organs, and were less abundant in esca and WD plants than in control plants (*P* = 10^-4^).

### Effects of drought and esca on wood integrity

We quantified the areas occupied by apparently healthy wood, black necrotic wood and white rot in trunk cross sections (Supplementary Table S7). The relative proportions of these three types of wood differed significantly between plant statuses (i.e. control, esca and WD; *P* = 10^-4^). The proportion of healthy wood was significantly lower in plants with esca symptoms than in control plants, but did not differ significantly between WD plants and the other two categories (Fig. 3A). The proportions of black necrosis and white rot did not differ significantly between plant disease statuses (*P* = 0.10 and *P* = 0.12, respectively), but there was a trend towards a higher proportion of white rot in plants with esca symptoms and a higher proportion of black necrosis in WD plants (Fig. 3A). In 60% of plants with esca symptoms, the proportion of healthy wood was lower than the minimum value recorded in control plants (34.3%).

**Fig. 3.**
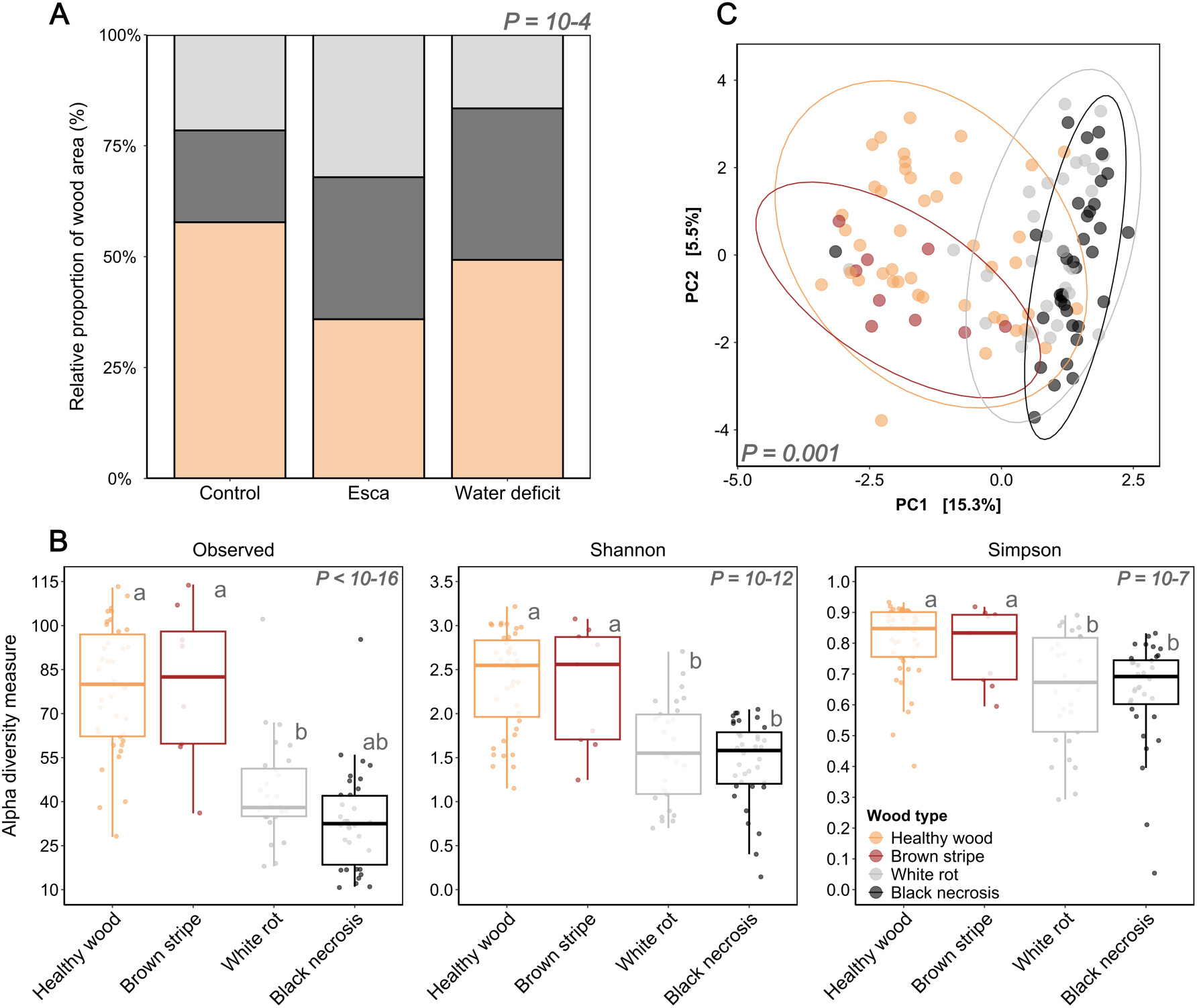
Relative proportions of wood symptoms in trunk cross-sections according to plant status (control, esca expression, water deficit), and the properties of the endophytic Ascomycota communities associated with each wood type. **A**, Mean proportions of healthy wood, black necrosis and white rot for each set of experimental conditions. **B**, Effect of wood type on the alpha-diversity metrics (i.e. observed richness, Shannon and Simpson indices) associated with Ascomycota communities. **C**, PCA summarizing the structure of Ascomycota communities according to an RDA on CLR-transformed data. Ellipses correspond to the 95% interval for each group. **D**, OTUs displaying significant differences in abundance between wood types according to a LeFse analysis. Features with an LDA score > 2 and a Kruskal-Wallis *p*-value < 0.05 are shown, and each bar is colored according to the group in which the OTU is overrepresented. **E**, Relative abundance of putative wood pathogens, putative antagonists and other taxa across wood types. All graphs are colored according to wood type.

### Ascomycota communities in trunk necrotic wood tissues

We compared trunk Ascomycota communities between apparently healthy wood, brown stripe, black necrosis and white rot samples collected in October, several weeks after symptom onset. Brown stripe samples were collected only from plants displaying esca symptoms (Table 1) and the sample size is lower for this tissue type. Wood type significantly affected alpha-diversity metrics, i.e. observed richness, and the Shannon and Simpson indices (*P* < 10^-16^, *P* = 10^-12^ and *P* = 10^-6^, respectively). Ascomycota richness and diversity were lower in necrotic wood (i.e. black necrosis or white rot), than in healthy wood and brown stripe (Fig. 3B). PCA and PERMANOVA analysis revealed that wood type accounted for 11.9% of the variance of the Ascomycota community in a model including plant status and wood type * plant status as covariates (*P* = 0.001). All pairwise comparisons between wood types were significant, but the clearest difference was that between necrotic and non-necrotic areas (Fig. 3C). The interaction between wood type and plant status was not significant: the differences in Ascomycota communities between non-necrotic and necrotic wood were observed in control, esca or WD plants (Supplementary Fig. S3).

Classifying Ascomycota OTUs into putative wood pathogens, antagonists and others (Supplementary Table S1), we found that wood type significantly modified the relative abundance of putative wood pathogens and putative antagonists, but not of other taxa (*P* = 10^-15^, *P* = 10^-10^ and *P* = 0.16, respectively; Fig. 4A). The abundance of putative pathogens was higher within the areas of black and white rot than in healthy wood (Fig. 4A). The abundance of putative antagonists was high in healthy wood and brown stripe, whereas these fungi were almost entirely absent from necrotic areas (Fig. 4A). Finally, we performed a linear discriminant analysis to identify the Ascomycota species driving the differences in microbial composition between wood types. Six fungal species, including the putative antagonist of trunk pathogens *E. nigrum,* were identified as biomarkers of healthy wood. Nine fungal species were identified as biomarkers of brown stripe samples, including several pathogens belonging to the Botryosphaeriaceae and putative antagonist *Cladosporium* spp. Three fungal species were significantly associated with black necrosis, including the pathogenic species *P. minimum*. Four fungal species were significantly associated with white rot, including the pathogenic species *P. chlamydospora* and several *Penicillium* spp. (Fig. 4B).

**Fig. 4.**
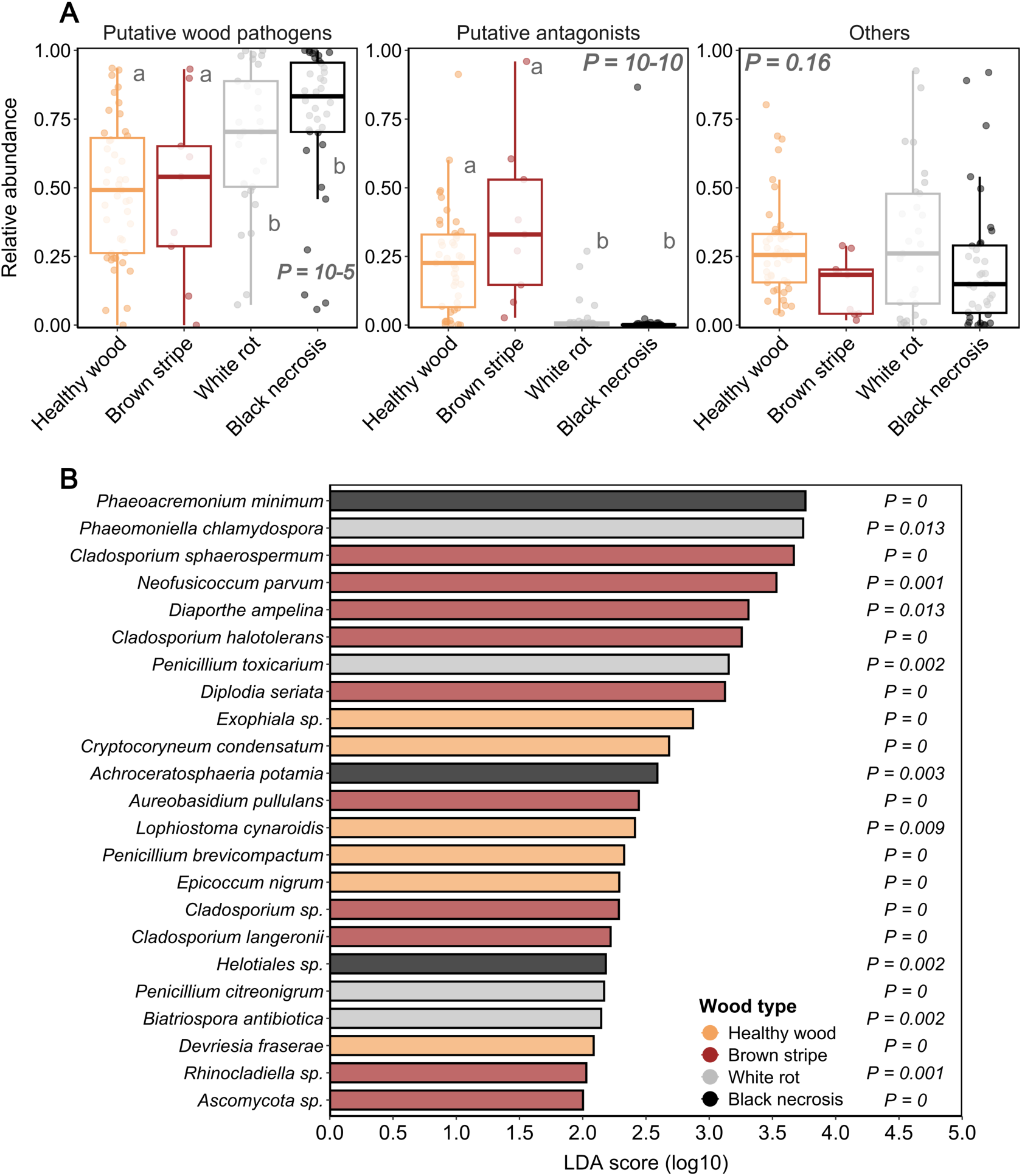
Functional and taxonomic characterization of Ascomycota communities across wood types. **A**, Relative abundance of putative wood pathogens, putative antagonists and other taxa, by wood type. All graphs are colored according to wood type. **B**, OTUs with significantly different abundances between wood types according to a LeFse analysis. Features with an LDA score > 2 and a Kruskal-Wallis *p*-value < 0.05 are shown, and each bar is colored according to the group in which the OTU is overrepresented.

### Effects of esca history on wood necrosis and Ascomycota communities

We assessed the effect of esca history on the distribution of wood types and the properties of Ascomycota communities in control plants. Esca history (comparing pS and pA plants with plant status as a covariable) had no significant impact on the distribution of wood types (healthy wood, black necrosis and white rot) in trunk cross sections (Supplementary Table S8).

In addition, the Ascomycota community in healthy trunk wood and young organ samples did not differ significantly between pA and pS plants in terms of either alpha-diversity or beta-diversity metrics (Supplementary Table S8).

## Discussion

Disentangling the complex interplay between plants, microbial communities and abiotic stresses is crucial for understanding the processes underlying perennial plant decline. In this study, we used a unique experimental design to characterize plant and Ascomycota community responses to esca disease and drought stress in mature grapevines. We extended the work of Bortolami et al. (2021b) showing that the expression of esca symptoms is inhibited by drought under controlled conditions, using naturally infected plants. We sampled the plants at the end of the experiment and demonstrated that the impact of drought on Ascomycota communities was greatest in the trunk, whereas the expression of esca symptoms affected mostly younger organs. The esca-drought antagonism revealed by Bortolami et al. (2021b) is not related to a decrease in the abundance of grapevine wood pathogens. Finally, we demonstrated that the incidence of esca symptoms on leaves was associated with a significant decrease in the relative proportion of apparently healthy (i.e. non-necrotic) wood in the trunk, with a corresponding increase in the proportion of necrotic tissues with a low diversity of Ascomycota and a high abundance of wood pathogens.

### Drought favors the pathogenic component of the Ascomycota community in trunk healthy wood whereas esca does not

In this study, the endophytic Ascomycota community in the healthy wood of the trunk was particularly strongly affected by intense drought stresses applied during two consecutive summers, whereas the seasonal expression of esca leaf symptoms affected the Ascomycota within young organs. The impact of drought has been studied in belowground communities in particular (e.g. Carbone et al. 2021, Jaeger et al. 2024), but drought can also shape endophytic microbial communities, by (i) directly affecting the living environment of fungi, by decreasing xylem water potential and humidity (Aung et al. 2018, Luard and Griffin 1981, Dell’Acqua et al., 2025) and (ii) modulating microbe-microbe and plant-microbe interactions (Trivedi et al. 2020). We found a low diversity of Ascomycota in healthy trunk wood under drought conditions. This finding is consistent with the results of Leal et al. (2024), who used one-year-old grapevine cuttings to demonstrate a substantial decrease in microbial interactions, thereby providing evidence in support of a drought-induced microbial imbalance. The effects of drought stress on plant and microbial communities include osmotic shock and the generation of reactive oxygen species (ROS; Schimel et al. 2007; Miller et al. 2010, Singh et al. 2011). Under drought conditions, the Ascomycota community in the trunk may shift towards specific taxa with more efficient intracellular ROS scavenging mechanisms and an ability to produce compatible osmolytes. The relative abundance of Ascomycota wood pathogens, including *P.chlamydospora* in particular, increased in the trunk under water deficit conditions, even in the absence of foliar symptoms on water-stressed plants (Bortolami et al. 2021b). The increase in *P. chlamydospora* levels under drought conditions has been reported before (Hrycan et al. 2025, Leal et al. 2024). It may be due to a change in fungal interactions, with an increase in the abundance of competitive species and their activities under drought, and a greater adaptation of *P. chlamydospora* to a constrained environment (i.e. in the presence of activated plant defenses and a low water potential) potentially favoring its development at the expense of its usual antagonists and competitors. Interestingly, *P. chlamydospora* seems to cope well with the oxidative stress induced by plant cells due to its efficient ROS scavenging arsenal (Fischer et al. 2016). However, this finding is more surprising given that the foliar expression of esca symptoms was inhibited under drought conditions.

During the years following rewatering, the WD plants kept alive had a probability of expressing esca leaf symptoms similar to that of control plants (C. Delmas, personal communication). The increase in *P. chlamydospora* abundance did not lead to a higher incidence of symptoms in subsequent years. The total wood pathogen load appeared to be higher in plants expressing esca, due to the presence of larger necrotic areas rich in pathogens. However, the observation that these pathogens are not more abundant in the healthy wood of symptomatic plants, where the process of degradation leading to necrosis presumably occurs, raises questions about the exact role of Ascomycota - accounting for around 80% of the whole fungal community in grapevine wood (Chambard et al. 2025, Jayawardena et al. 2018) - in esca pathogenesis. Ascomycota pathogens are generally considered to play an important, if not essential role in the expression of esca symptoms (e.g. Bertsch et al. 2013, Gramaje et al. 2018). However, relative abundance, studied here through metabarcoding approaches, may not be the most relevant trait. Instead, the activity of Ascomycota pathogens (e.g. *P. chlamydospora*, *Botryosphaeriaceae* spp.), including their expression of virulence factors, may decrease in water deficit conditions, or increase before the expression of esca symptoms (Desprez-Loustau et al. 2006, Morales-Cruz et al. 2018a, Nerva et al. 2022). A recent study on the same plants as the present one did not confirm these hypotheses. Chambard et al. (2025) highlighted an increase of *P. chlamydospora* activity (growth, reproduction) under drought, and no modification of Ascomycota wood pathogen behaviour following esca symptom expression. Thus, there is growing scientific evidence to suggest that the contribution of Ascomycota to esca expression is overestimated (Hofstetter et al. 2012, Nerva et al. 2022, Moretti et al. 2023, Chambard et al. 2025). These results call for refining our knowledge of gene expression and metabolic changes in both plants and pathogens in the context of multiple stress experiments.

Drought did not significantly modify the proportion of necrotic wood, but it did slightly decrease the white rot/black necrosis ratio. This suggests that drought may interfere slightly with the activity of white-rot basidiomycetes that degrade wood through both enzymatic and non-enzymatic decay pathways (Moretti et al. 2021, 2023). Drought has already been identified as a variable that may affect wood decay (Jia et al. 2024). Indeed, to grow through the lignocellulose matrix and release their arsenal of decay-mediating molecules (i.e. enzymes and low-molecular weight compounds) into the wood cell wall, wood decay fungi (Basidiomycota included) require the moisture content in the wood to be at least above the fiber saturation point (Goodell 2020, Zaber and Morrell 2012). This highlights the importance of studying the abundance and activity of Basidiomycota, but also of bacteria that can act in synergy in the wood degradation process (Del Frari et al. 2021, Haidar et al. 2021).

The expression of esca symptoms did not significantly modify plant-microbiome interactions in healthy trunk tissues (despite a trend towards lower species richness and an increase in the abundance of wood pathogens) as reported in other studies (Del Frari et al. 2019, Monod et al. 2024; Dell’Acqua et al. 2025). Very few studies have quantified the impact of dieback diseases on endophytic communities in other plants (e.g. Raghavendra et al. 2017, Steinrucken et al. 2016), although modifications in belowground microbial diversity were highlighted (Darriaut et al. 2024, Farooq et al. 2022, Han et al. 2022, Lasa et al. 2024). Such modifications may be accompanied by an ecological disequilibrium, with an increase in the pathogenic component and a loss of beneficial functions, such as symbioses between plants and fungi and the promotion of plant growth (Gómez-Aparicio et al. 2022, Ma et al. 2024, Solís-García et al. 2021). Our findings confirm that esca is a complex pathosystem, in which plant physiology, interacting with the abiotic environment, plays a key role in leaf symptom onset, without any major change in fungal communities in healthy wood tissue.

### Esca induces a more significant shift in Ascomycota communities in young organs than drought

Two consecutive summer seasons under drought conditions had a greater impact on the trunk than esca, but seasonal esca expression had a greater effect on Ascomycota communities in younger organs, which undergo significant physical, physiological and metabolic changes during the expression of esca symptoms (Bortolami et al. 2021a,b, Dell’Acqua et al. 2024, 2025). The Ascomycota communities inhabiting young organs (mainly endophytes, except for the petioles, from which we sampled both endophytes and epiphytes) on plants displaying esca symptoms were less diverse than those of control plants and were largely dominated by *Cladosporium* species. This finding suggests that the presence of leaf symptoms influences, albeit slightly, microbial communities distal to the infection zone characterized by necrosis and abundant trunk pathogens (i.e. perennial organs). Leaf endophytic fungal communities have been shown to be modified by esca expression (Del Frari et al. 2025). Accumulation and qualitative changes have been reported for primary and secondary metabolites (e.g. sugars, terpenes, flavonoids; Bortolami et al. 2021b, Del Frari et al. 2025, Weiller et al. 2024) in leaves and stems expressing esca symptoms, and these effects may be greater than those observed in the trunk, explaining the stronger response of fungal communities to esca expression in younger organs. Moreover, biotic stresses (such as esca disease) also induce ROS generation, and fungi have developed strategies for scavenging ROS (Lehmann et al. 2015). Therefore, as discussed above for the trunk, Ascomycota communities in young organs may shift towards specific taxa with more efficient ROS scavenging mechanisms.

However, esca had only a small effect in our experiment and other studies have revealed the stem microbial communities to be similar in asymptomatic plants and plants with esca symptoms (Del Frari et al. 2019). Water deficit had an even weaker effect on the Ascomycota communities of young organs, despite the highly significant physiological impact of drought during the experiment (Bortolami et al. 2021b). The main physiological difference between the plants subjected to water deficit and those expressing esca symptoms was that water potential was strongly affected by drought but not by esca expression, despite both conditions decreasing plant transpiration. Photosynthesis and the balance of non-structural carbohydrates were more strongly affected by esca expression, with a decrease in leaf chlorophyll content and an accumulation of monosaccharides in leaves and stems, at the expense of sucrose production and starch storage (Bortolami et al. 2021b). The Ascomycota communities in young organs may have been more sensitive to carbon resource availability and the concentration of non-structural carbohydrates, whereas wood water status may be ecologically more selective for trunk communities. Conversely, these results demonstrate the resilience of the microbial communities in young aerial organs to decreases in water potential during drought, with no significant decrease in diversity and no increase in the abundance of specific ecological guilds. Finally, the relative abundance of *E. necator*, the causal agent of grapevine powdery mildew, was higher in asymptomatic vegetative organs on plants displaying esca symptoms, potentially due to a slight increase in gas exchange in these specific organs (Bortolami et al. 2021b). *E. necator* was less prevalent in organs expressing esca symptoms, perhaps due to the impairment of carbon assimilation (Guilpart et al. 2017).

### On the relationship between wood necrosis, pathogens and the expression of esca foliar symptoms

The causal links between the internal consequences of plant-pathogen interactions (i.e. wood necrosis) and the external expression of disease (i.e. foliar symptoms) remain poorly understood in perennial plant decline (Claverie et al. 2020, Gosling et al. 2024, Skovsgaard et al. 2010). Esca foliar symptoms have been reported to be associated with a higher proportion of wood necrosis, especially white rot, in *Vitis vinifera* L. cv. ‘Cabernet-Sauvignon’ (Maher et al. 2012, Ouadi et al. 2019). Consistent with these findings, we found that plants displaying esca symptoms had significantly lower proportions of healthy wood in their trunks than control plants. However, our results demonstrate that the proportion of healthy wood explained esca symptom expression in *V. vinifera* cv. ‘Sauvignon blanc’ better than the type of necrosis. This may reflect a cumulative loss of functional xylem area (i.e. of the active tissues located in the outer xylem; McElrone et al. 2021) in symptomatic plants, regardless of the type of degradation. The mean proportion of white rot was about 20% in control plants (i.e. well-watered and asymptomatic in the year of sampling), higher than in previous reports. As plant age and genotype are significant drivers of esca leaf symptom expression (Etienne et al. 2024, Gastou et al. 2024), we can hypothesize that these factors modulate the proportion of necrotic wood (Claverie et al. 2023). In old plants (30-y-o in our study), control plants with an asymptomatic history had similar necrosis extent and Ascomycota community in the trunk than previously symptomatic plants. In addition to a threshold of necrotic wood proportion in the trunk, the area of contact between active xylem tissue (efficiently transporting sap) and specific necrotic tissues, or the activity of microbial communities (e.g. virulence factors) may be involved in the onset of foliar symptoms and should be explored further.

In the classical model of plant-microorganism interactions underpinning esca expression, Ascomycota fungi are considered to be responsible for black necrosis, black spots, wood streaks and discoloration, whereas white rot is associated with Basidiomycota wood-degrading species, probably in interaction with bacteria (Bertsch et al. 2013, Bruez et al. 2020, Haidar et al. 2021, 2024, Moretti et al. 2021). However, less attention has been paid to the relationship between the total microbiome and the type and extent of necrosis and to the role of Ascomycota fungi in white rot propagation. We show here that, whatever the health status of the plant (i.e. control, Esca or water deficit), the necrotic areas in the trunk harbored less diverse Ascomycota fungal communities than adjacent apparently healthy areas, confirming the results obtained for cordons (Bruez et al. 2020). These areas were dominated by Ascomycota wood pathogens, the proportion of which increased at the expense of other taxa, including putative antagonists, as previously highlighted in grapevine trunk diseases (Martín et al. 2022), but not in other pathosystems (Bakys et al. 2009, Denman et al. 2016). We found that *P. minimum* was clearly overrepresented in black necrotic areas, whereas *P. chlamydospora* was not. *P. chlamydospora* was significantly overrepresented in white rot tissues, as previously reported by Bruez et al. (2020) and Pacetti et al. (2021). The previously favored hypothesis of a pioneering role of Ascomycota fungi before Basidiomycota species in the sequence of events occurring during the colonization of healthy grapevine wood has recently been revisited in light of new findings on potential intra- and extracellular machinery for the detoxification of wood phenolic compounds in *F. mediterranea* (Moretti et al. 2021, Schilling et al. 2022, Garcia et al. 2024). For *P. chlamydospora* a hypothetical role in wood decay through the opportunistic colonization of highly degraded areas or co-degradation, possibly alongside Basidiomycota, therefore appears more plausible than the pioneer hypothesis. Moreover, *P. chlamydospora* may be less closely associated with black necrosis, as previously suggested (e.g. Bertsch et al. 2013, Surico et al. 2006). It is, in any case, a very abundant endophyte in healthy tissues (Del Frari et al. 2019, Hofstetter et al. 2012, Dell’Acqua et al. 2025). Finally, we resequenced healthy wood samples, using ITS markers to quantify Basidiomycota and determine their relative abundance in healthy wood samples; we confirmed that *F. mediterranea* was present at high abundance in the wood of plants expressing esca symptoms (Chambard et al. 2025). The activity of *F. mediterranea* was also significantly altered, with overexpression of genes involved in growth, nutrition, and interactions with the plant and other fungal species (Chambard et al. 2025). While the co-occurrence of white-rot and esca expression is established (Maher et al. 2012, Ouadi et al. 2019), a direct causal link between the abundance and activity of Basidiomycota pathogens in wood and the development of external symptoms has yet to be demonstrated.

We also characterized the Ascomycota community associated with the brown stripe xylem sampled about three months after the onset of leaf symptoms. We showed that the Ascomycota community in this symptomatic tissue is diverse and similar to that retrieved from healthy woody tissues. This finding is consistent with the fact that the brown stripe has not yet become a necrotic area. This tissue no longer conducts water, as shown by the use of a dye to follow sapwood transport (Lecomte et al. 2024), and in other vascular pathosystems (Agrios 2005, Bibuci and Cirulli, 2012, Jiménez-Díaz et al. 2012). The brown color probably results from the translocation of oxidizing phytotoxins and degradation products and is associated with non-gaseous vascular embolism (e.g. tyloses and gels; Agrios 2005, Bubici and Cirulli, 2012, Lecomte et al. 2024). We confirmed the high abundance of Botryosphaeriaceae spp. in brown stripe tissues late in the season (Larignon et al. 2001), alongside *Cladosporium* spp. and *A. pullulans*. The latter was reported in the literature as an antagonist of wood pathogens (Mesguida et al. 2023) and, in our experiment, was particularly abundant in control vines compared to stressed ones. Conversely, *in planta*, some strains are associated with esca leaf symptom severity, and may act synergistically with *P. chlamydospora* to induce wilts (Karácsony et al. 2023). Hence, its role in esca pathogenesis deserves further investigation, to determine its contribution (positive or negative) to internal and external symptom expression, and its activity within brown stripes. More generally, observations were performed three months after symptom onset. We cannot therefore draw any firm conclusions about the changes in plant-fungus interactions potentially leading to the co-occurrence of brown stripe and foliar symptoms. Detailed studies of the microbial and physiological features related to the brown stripe symptom, and of the temporal dynamics linking internal and external symptoms, are now required.

## Conclusion

Overall, this experimental study provides a new perspective on the ways in which abiotic and biotic stresses can shape microbial communities in perennial plants and contribute to plant decline. The trunk and younger grapevine organs had different endophytic fungal communities with different responses to drought and esca. Only esca significantly altered wood integrity. However, we were unable to investigate the dual-stress situation, as water deficit inhibited the expression of esca foliar symptoms. Based on these findings, assessments of the responses of the grapevine holobiont to combinations of stresses at the physiological and microbiome levels across various cultivars appear relevant to identify promising genotypes in the context of climate change and perennial plant decline. Indeed, wood degradation (Arnold et al. 2025), the incidence of esca foliar symptoms (Gastou et al. 2024), drought tolerance (Dayer et al. 2022) and endophytic microbial communities (Bekris et al. 2021, Hamaoka et al. 2022) differ between grapevine genotypes. Finally, plant dieback is related not only to endophytic communities, but also to microbiomes inhabiting the soil compartment (Darriaut et al. 2021, 2023, Gómez-Aparicio et al. 2022, Lasa et al. 2024). Future holistic *in natura* microbiome studies (i.e. multidisciplinary, multicompartments and/or multitaxa) would be particularly helpful, providing precise and generalizable results on the response of the holobiont to stressful conditions.

## Data availability

DNA sequencing datasets are available in the ENA database under the accession number PRJEB87388.

## Acknowledgments

We would like to thank Elena Farolfi and Jérôme Pouzoulet for help with plant sectioning and wood sampling, and Corinne Vacher for advice on metabarcoding analyses.

## Author contributions

C.E.L.D., G.A.G. and G.B. conceived the experimental drought x esca study; G.B. and N.F. sampled the tissues and ground the samples; G.B. analyzed the trunk section images; N.F. extracted DNA and performed PCR; P.G and J.V processed the bioinformatics pipeline; P.G. analyzed the data; S.M. contributed to the discussion section and to data interpretation; P.G wrote the first draft of the paper under the supervision of C.E.L.D.; All authors critically revised and approved the final version of the manuscript for submission.

## List of supplementary material

**Supplementary Fig. S1:** Pictures of the sampling

**Supplementary Fig. S2:** Properties of Ascomycota communities in all samples (i.e. trunk, cane, stem and petiole samples) across organs.

**Supplementary Fig. S3:** Ascomycota community structure across wood types for each plant status.

**Supplementary Fig.S2:** Properties of Ascomycota communities in all samples (i.e. trunk, cane, stem and petiole samples) across organs.

**Supplementary Table S1:** Ascomycota genera classified as putative wood pathogens and putative wood antagonists.

**Supplementary Table S2:** Global metrics describing the GTAA metabarcoding dataset before and after the decontamination process.

**Supplementary Table S3:** OTU table corresponding to the entire decontaminated GTAA metabarcoding dataset.

**Supplementary Table S4:** List of experimental factors and Ascomycota reads for each sample included in the GTAA metabarcoding study.

**Supplementary Table S5:** Summary of the results obtained in Aldex2 differential analyses in healthy trunk samples and young samples.

**Supplementary Table S6:** Statistical modelling of organ health status (S-AS vs S-S) effects on alpha-diversity metrics and Ascomycota community structure computed for young organs.

**Supplementary Table S7:** List of experimental factors and wood type percentages for each plant

**Supplementary Table S8:** Statistical modelling of esca history (pA vs pS) effects on the distribution of wood types and alpha-diversity metrics and Ascomycota community structure computed for healthy trunk and young samples.

