## Supplementary materials for "Differential impacts of drought and esca expression on Ascomycota fungi in the trunks and young organs of mature grapevines"

### Supplementary figures

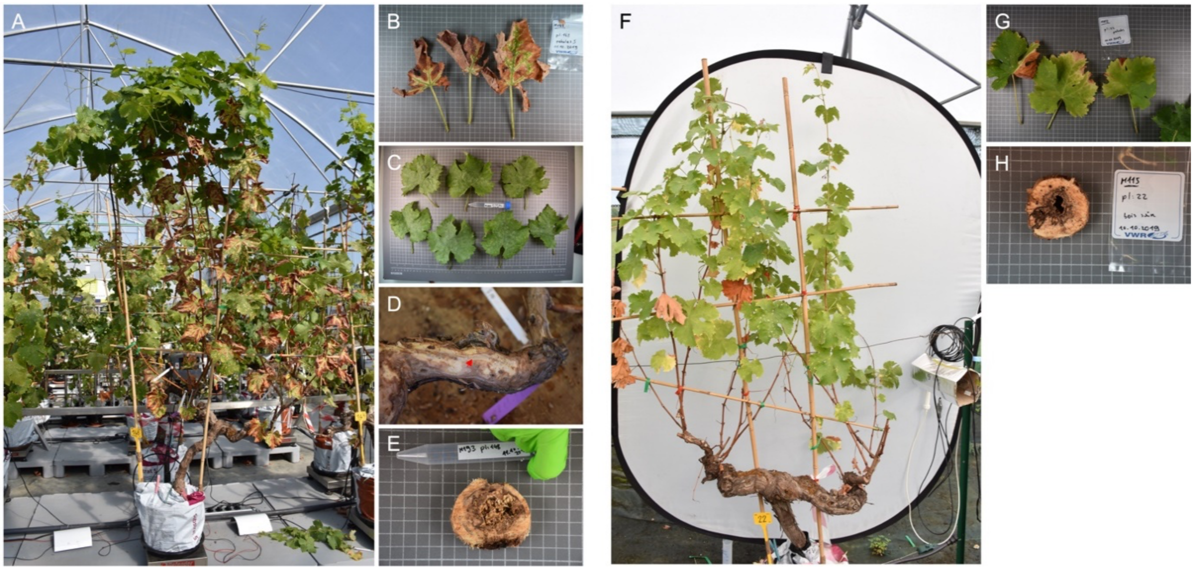


**Supplementary Fig. S1: Pictures of the sampling**. **A**, *Vitis vinifera* cv. Sauvignon blanc expressing esca leaf symptoms, from which we sampled petioles from **B** symptomatic or **C** asymptomatic leaves, **D** wood tissues from the xylem brown stripe (red arrow), different wood tissues collected from **E** a trunk cross section, as well as one-year-old (stems) and two-year-old (canes) shoots (not picturised). **F**, *Vitis vinifera* cv. Sauvignon blanc under water deficit, from which we sampled petioles from **G** leaves, different wood tissues collected from **H** a trunk cross section, as well as one-year-old (stems) and two-year-old (canes) shoots (not picturised). Similarly, we sampled the same types of tissue from control well-watered asymptomatic plants (not shown).

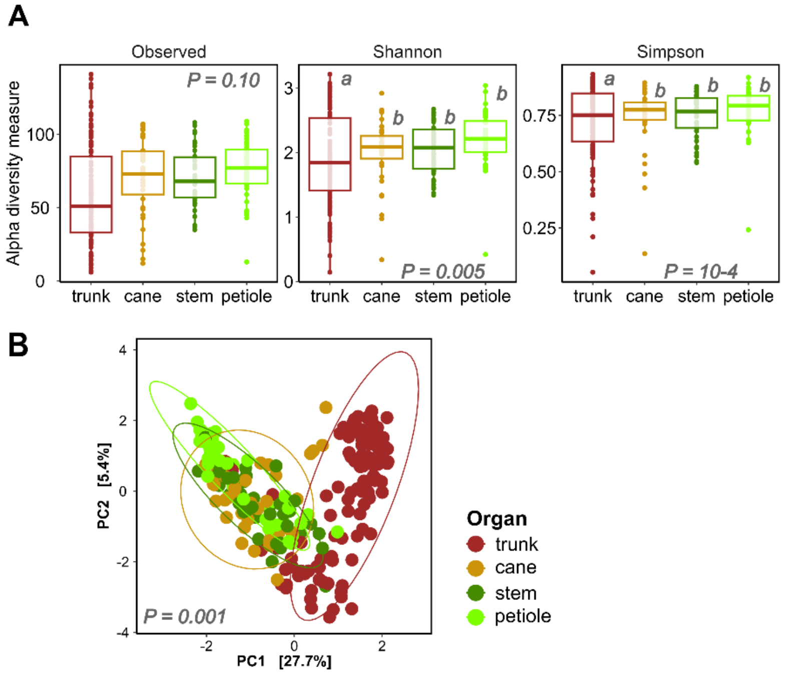


**Supplementary Fig. S2: Properties of Ascomycota communities in all samples (i.e. trunk, cane, stem and petiole samples) across organs. A**, Effect of the organ on alpha-diversity metrics (i.e. observed richness, Shannon and Simpson indexes) associated with Ascomycota communities. P-values are calculated in ANOVA procedures, and letters correspond to the groups of significance according to Tukey tests with an alpha risk of 5 %. **B**, PCA summarising the structure of Ascomycota communities according to a RDA on CLR-transformed data. P-value is calculated in a PERMANOVA procedure, ellipses correspond to the 95% interval for each group. *All graphs are coloured according to the organ. Cane, stem and petiole samples will subsequently be referred as « young » organs.*

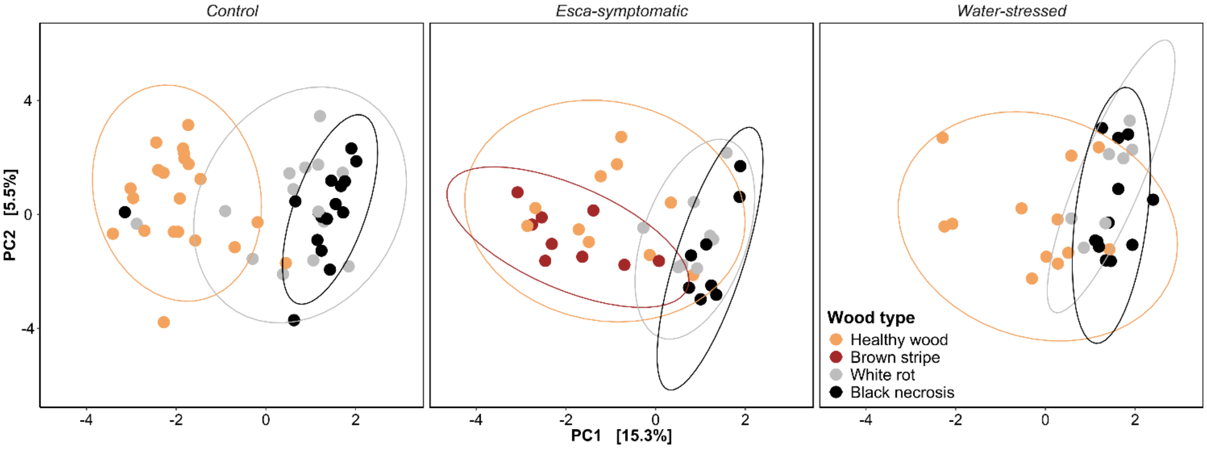


**Supplementary Fig. S3: Ascomycota community structure across wood types for each plant status.** PCA summarising the structure of Ascomycota communities according to a RDA on CLR-transformed data. Ellipses correspond to the 95% interval for each group. *All graphs are coloured according to wood type.*
